# Enteroaggregative *Escherichia* clade I from Nigeria

**DOI:** 10.64898/2026.04.21.719883

**Authors:** Rotimi A. Dada, Olabisi C. Akinlabi, Babajide A. Tytler, Busayo O. Olayinka, Andrew J. Page, Nicholas R. Thomson, Iruka N. Okeke

**Affiliations:** Department of Pharmaceutical Microbiology and Biotechnology, Faculty of Pharmaceutical Sciences, Ahmadu Bello University, Zaria, Nigeria; Department of Pharmaceutical Microbiology, Faculty of Pharmacy, University of Ibadan, Ibadan, Nigeria; Origin Sciences, Cambridge, United Kingdom; Parasites and Microbes Programme, Wellcome Sanger Institute, Cambridge, United Kingdom; London School of Hygiene and Tropical Medicine, London, United Kingdom

## Abstract

*Escherichia coli*, the *Escherichia* type species, is present in mammalian and avian intestinal microbiota, and includes both commensals and pathogens. Other *Escherichia* species are understudied because they are less commonly associated with human disease and because of paucity of tools that can correctly delineate them from *E. coli*. However, other species of this genus including *Escherichia albertii* and *Escherichia fergusonii* are repeatedly reported as diarrhoeagenic. We hypothesized that some bacteria fitting the definition of enteroaggregative *E. coli* (EAEC) belong to species other than *E. coli*. We used phylogeny to determine the species of 2,818 *Escherichia* genomes from diarrhoea epidemiology studies in Nigeria. Phylogeny speciation was confirmed using GTDB-tk and ClermonTyping. Virulence genes were detected using ARIBA/Virulencefinder database and multilocus sequence typing performed using the Achtman scheme. Fourteen non-coli *Escherichia* genomes were identified— *Escherichia* clade I ST485 (11), *Escherichia ruysiae* ST5792 (2) and *Escherichia fergusonii* ST5636 (1). All the *Escherichia* clade I ST485 carry EAEC virulence genes *aap, aar, astA* and *air,* as well as *hlyF, eatA, tsh*, *traT,* and *chuA* virulence genes. Interestingly, 62% of enteroaggregative *Escherichia* clade I ST485 genomes listed on Enterobase are from Africa isolates, despite only 3% of genomes overall coming from the continent. Our results suggest that non-*coli Escherichia* species are infrequently isolated from human stool, but, when they are, they are misidentified as *E*. *coli* so that their significance is largely overlooked. *Escherichia* clade I ST485 is a globally disseminated enteroaggregative *Escherichia* clade I lineage that is common in Africa.

**Author Summary:** Escherichia clade I are rarely associated with disease and because of the difficulty in differentiating them from *Escherichia coli* in routine laboratory, they are often misidentified as *Escherichia coli* leading to the underestimation of their impact on the burden of disease. Additionally, some clones of *Escherichia* clade I also carry genetic markers that have been used to define Enteroaggregative *Escherichia coli* (EAEC), a cause of persistent diarrhoea in developing countries and traveller’s diarrhoea in developed economies. EAEC has also been associated with malnutrition and poor growth among children in developing economies. We here describe clones of *Escherichia* clade I (ST485) that carries enteroaggregative genes and in some cases, recovered from diarrhoeal cases. We show from genomes deposited on Enterobase and our study, that this clone is globally disseminated, often associated with human infections and often misidentified as *Escherichia coli*. We also describe other non-*coli Escherichia* other than *Escherichia* clade I isolated from humans. We suggest that the *Escherichia* clade I clone carrying enteroaggregative genes may be described as Enteroaggregative *Escherichia* clade I.

## Introduction

The genus *Escherichia* encompasses considerable diversity [1,2,3,4,5,6,7]. *Escherichia* species other than *E. coli* were originally recognized as numbered ‘clades’. While *Escherichia* clade I, the most closely related to *Escherichia coli*, still retains its ‘clade’ nomenclature [8], other clades have received species names [3,8,9,10]. *Escherichia* clade II is now referred to as *Escherichia whittamii* (*E*. *whittamii*), clades III and IV are now *Escherichia ruysiae* (*E*. *ruysiae*) and clade V, *Escherichia marmotae* (*E*. *marmotae*).

*Escherichia coli* is a commensal found in the gut of humans, other mammals and birds. Most *E. coli* are harmless and are thought to be involved in maintaining the balance of the gut microbiota. However, diarrhoeagenic *E. coli* (DEC) and extraintestinal pathogenic *E. coli* (ExPEC) can cause intestinal and extraintestinal disease, respectively. *Shigella* ‘species’, historically thought to be distinct from *Escherichia* because they cause dysentery and are less biochemically active, represent multiple *E. coli* lineages that have convergently gained virulence and lost housekeeping genes through negative selection [11,12,13,14]. Within the *E. coli* species, different DEC and ExPEC pathotypes, including *Shigella,* have specific virulence gene repertoires and pathogenic features. In some cases, pathotypes are lineage-derived and in others they share mobile genes or pathogenic phenotypes.

A few reports have implicated strains of non-*coli Escherichia* in human disease. In Australia, [15] reported the isolation of *E. marmotae* from a urine sample. Initial and repeat samples of that bacterium were phenotypically identified as *E. coli*, including using the Vitek2 system (XL GNI ID cards), which gave the *E. coli* identification with 99% probability. However, MALDI-TOF-MS identification returned the identity of the strain as *E. marmotae* with a confidence score of 2.39 and 16S rDNA sequencing showed 98.9% identity three known *E. marmotae* strains. In the Netherlands, *E. ruysiae* has been isolated from the stool of a returning traveller with diarrhoea [16] and was initially identified as ESBL-producing *E. coli*. A multidrug resistant strain of *E*. *fergusonii* reported in Italy was implicated as the cause of cystitis in a 52-year-old woman [17] and in China, [18] also reported a case of a 72-year-old male who developed skin and sclera yellow staining attributed to an infecting, biofilm-forming *E*. *fergusonii* strain. *E*. *fergusonii* have been repeatedly implicated in human diarrhoea and one strain was recently shown to carry an enterotoxigenic *E*. *coli* heat-labile enterotoxin 1 gene (*elt1*) on its chromosome [8]. *E. albertii* is even more frequently described and studied as a diarrhoeal pathogen, with multiple lineages carrying locus of enterocyte effacement pathogenicity islands, originally described in enteropathogenic and enterohemorrhagic *E. coli*., as well as cytolethal distending toxin genes [19]. Non-*coli Escherichia* have also been implicated in animal infections [20,21].

It is probable that non-*coli Escherichia* pathogens are more common but undetected or wrongly believed to be *Escherichia coli* because of lack of motivation and tools to delineate non-*coli Escherichia* from the *Escherichia coli*, especially in routine diagnostic microbiology laboratories [8,19]. The liberalisation and democratisation of genome sequencing coupled with availability of better speciation tools now makes it possible to rigorously re-interrogate presumptive *E. coli* pathotypes to determine whether they include non-*coli* species [22]. We have noticed that phylogenies of enteroaggregative *E. coli* (EAEC) from epidemiological studies in Nigeria typically include a few isolates external to the principal *E*. *coli* phylogroups. We therefore hypothesised that that some ‘*E. coli’* that fit the EAEC definition are members of other species. The aim of this study was to identify, determine the species and characterize putative non-*coli* enteroaggregative *Escherichia* species from diarrheal studies in Nigeria.

## Methods

We have heretofore sequenced a total of 3,320 faecal *Escherichia* genomes from five epidemiological studies: [23,24,25,26,27,28,29]. Post-Sequencing quality control (QC) was first conducted with FastQC [30] and the results were aggregated with MultiQC [31]. Kraken 2 [32] was used to assign genomic IDs to reads and Bracken [33] was used to quantify the genomic IDs assigned by Kraken. We used the genomic IDs assigned by Kraken/Bracken to set a cut-off point of ≥50% genomic markers for *Escherichia*-*Shigella* as inclusion criteria for genomes to proceed to downstream analyses. The genomes were mapped with EAEC 042 (FN554766.1) using a Wellcome Sanger Institute, Pathogen Informatics in-house tool—sh16_scripts (multiple_mappings_to_bam.py)—available here [34] producing a multifasta alignment file. SNPs were called out of the multifasta alignment files using snp-sites [35] and the SNPs were used to construct phylogenies using IQTree [36]. To provide some context, genomes of *E. coli* K12, *E. albertii, E. ruysiae, E. marmotae, E. whittamii, E. hermannii, E. vulneris, E. fergusonni* and *Salmonella enterica* serovariety Typhimurium were added to each of four sets of genomes. Each tree was rooted with *Salmonella enterica* serovar Typhimurium (SRR7426992). *Escherichia adecarboxylata* (now known as *Leclercia adecarboxylata*), and *Escherichia blattae* (now known as *Shimellia blattae*), which have been reclassified into other genera were excluded. The trees were viewed with Figtree [37], Phandango [38], iTOL [39] and Microreact [40]. After viewing the phylogenies, genomes representative of each lineage within *E. coli* phylogroups A, B1, B2, C, D, E, F and G, and all lineages that clustered outside of the *Escherichia* coli genus from each tree were combined to make a final tree. The final tree had genomes of *Escherichia* clade I, *E. albertii, E. ruysiae, E. marmotae, E. whittamii, E. hermannii, E. vulneris, E. fergusonni, Shigella dysentriae, Shigella boydii, Shigella flexneri, Shigella sonnei* and *Salmonella enterica* Typhimurium. For the final tree, TVM+F+ASC+R4 model was used and -bb 1000 option of IQTree was used to determine the Ultra-Fast Boostrap support values for 1000 replicates. The Shimondaira and Hasegawa approximate Likelihood-Ratio Test (SH-aLRT) [41], was also carried out using the -alrt 1000 option of IQTree. This was done to determine the log ratio between the likelihood value of the current tree and that of the best alternative. The final tree was viewed with Figtree [37], Phandango [38], iTOL [39] and Microreact [40]. Strains possessing of any of the following 13 virulence-associated genes (*aggR*, *aaiC*, *aap*, *aar*, *aatA*, *capU*, *air*, *eliA*, *aggA*, *aafA*, *agg3A*, *agg4A* and *agg5A*), which have previously been used to define EAEC [26,42,43,44,45,46,47,48,49] were designated enteroaggregative.

The Sequence Types (STs) of the genomes were determined using ARIBA [50] and MLST (Achtman 7 loci) database [51](https://pubmlst.org/). Virulence genes were detected using ARIBA [50] and VirulenceFinder database [52]. Resistance genes were identified using ARIBA and Resfinder database [53] and plasmid replicons were determined using ARIBA and Plasmidfinder database [54]. In-silico pathotyping was carried out using ARIBA [50] and VirulenceFinder database [52].

Sequence reads were assembled using SPAdes genome assembler [55]. Following sequence assembly, contigs generated by the SPAdes [55] were checked for quality of genome assembly using CheckM [56]. In-silico serotyping and phylogrouping were carried out using EcTyper [57] and Ezclermont [58] respectively.

The suspected non-*coli Escherichia* genomes were analysed using Genome Taxonomy Database-toolkit (GTDB-tk) [59] and ClermonTyping [60]. While GTDB-Tk is a toolkit for assigning taxonomic identification to bacterial and archaeal genomes, ClermonTyping tool is based on a concept of quadruplex PCR developed by Clermont and his colleagues. ANI, GTDB-tk and ClermonTyping speciation tests were performed, comparing each genome with those of the species node inferred from the phylogeny that we generated. For ANI values, all the *Escherichia* clade I were compared with *Escherichia coli*. *Escherichia fergusonii* with reference *Escherichia fergusonii* and *Escherichia ruysiae* with reference *Escherichia ruysiae*.

### Biochemical profiling

Isolates were maintained at -80°C in Luria Broth: Glycerol 1:1 and cultured on Tryptone Soya Agar (TSA) ahead of biochemical testing. 2-3 pure colonies from the TSA suspended in normal saline. The saline suspension was used for biochemical profiling on the Microbact 24E GNB in accordance with manufacturer’s directions (www.oxoid.com/UK/blue/prod_detail/prod_detail.asp?pr=MB1130). Microbact octal codes entered into software which gave the Microbact ID and % probability and individual results were used to study and compare profiles.

## Results

A total of 2,850 out of 3,320 whole genome sequences from isolates derived from five epidemiological studies previously described met the inclusion criteria for the study and 502 were excluded due to low abundance *E. coli* genomic markers. Based on phylogeny, 14 genomes that fell on branches linked to the *Escherichia* species reference genomes were identified as non-*coli Escherichia* and selected from the four phylogenetic trees. These are *Escherichia* clade I (11/14, 78.6%), *Escherichia ruysiae* (2/14, 14.3%) and *Escherichia fergusonii* (1/14, 7.1%) (Figure 1). No genomes falling within the *E*. *albertii, E*. *vulneris*, *E*. *marmotae* or *E*. *hermannii* species were identified.

**Figure 1:**
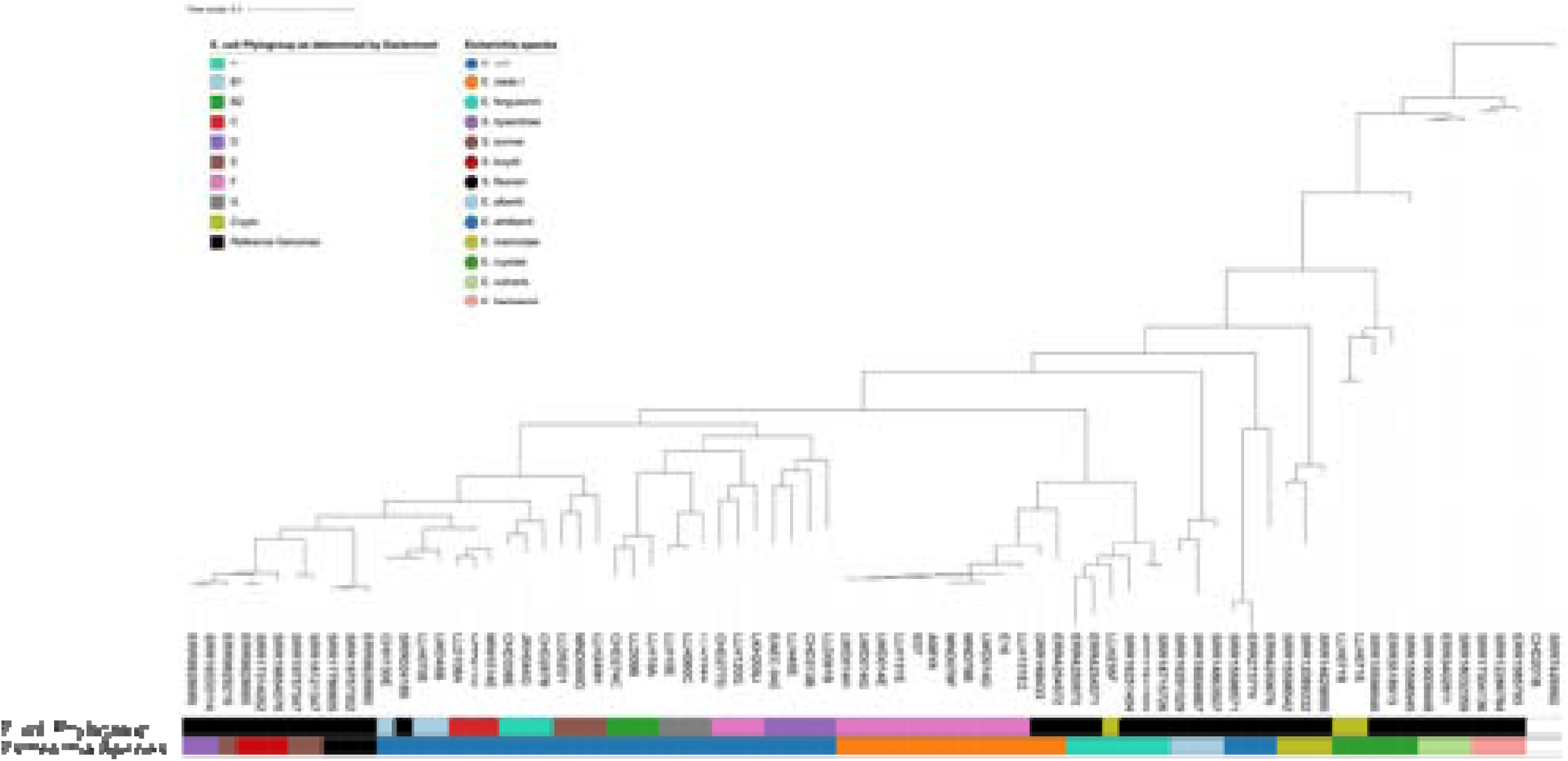
A maximum-likelihood phylogenetic tree that includes representatives of non *coli Escherichia* species, *E. coli* phylogenetic groups and *Shigella* serovarieties. Included in this genome set are typical and ‘atypical’ *E. coli* phylogenetic group F strains, the latter clustering with reference Escherichia clade I

Figure 1 shows a phylogenetic tree of 14 non-coli *Escherichia* with representative *E. coli* form phylogenetic groups A, B1, B2, C, D, E, F and G, *Shigella* species and reference non-*coli Escherichia* genomes. Each species clustered with other species in their lineages.

Of the 14 non-*coli Escherichia* genomes from stool samples, eight were from children with diarrhoea, five were from stools of apparently healthy children and one strain, A98YA, was from an adult with diarrhoea. Strains E07 and E18 were isolated in the mid 1990s, whilst the other isolates were recovered between 2015 and 2021.

As shown in Table 1, for species identification of *E*. *fergusonii* and *E*. *ruysiae*, the results of GTDB-tk tool aligned with the phylogeny generated using IQTree. However, this was not the case for other non-*coli* species. All the *Escherichia* clade I (from our phylogeny) were identified as *E. coli* by the GTDB-tk tool. ClemonTyping on the other hand, identified all *Escherichia* clade I and *Escherichia fergusonii* by their species names and *E*. *ruysiae* by its old name, *Escherichia* clade IV. Table 1 also shows the phylogeny-based identification of the non-*coli Escherichia*, their phylogenetic groups, ST and loci used for assigning their ST. Ezclermont *E*. *coli* phylogrouping tool assigned all enteroaggregative *Escherichia* clade I genomes to phylogenetic group F. *Escherichia fergusonii* and *E*. *ruysiae* genomes were assigned cryptic/undefined phylogenetic group. For the MLST, all *Escherichia* clade I were assigned to ST485 by ARIBA (Achtman 7-loci MLST scheme).

**Table 1:**
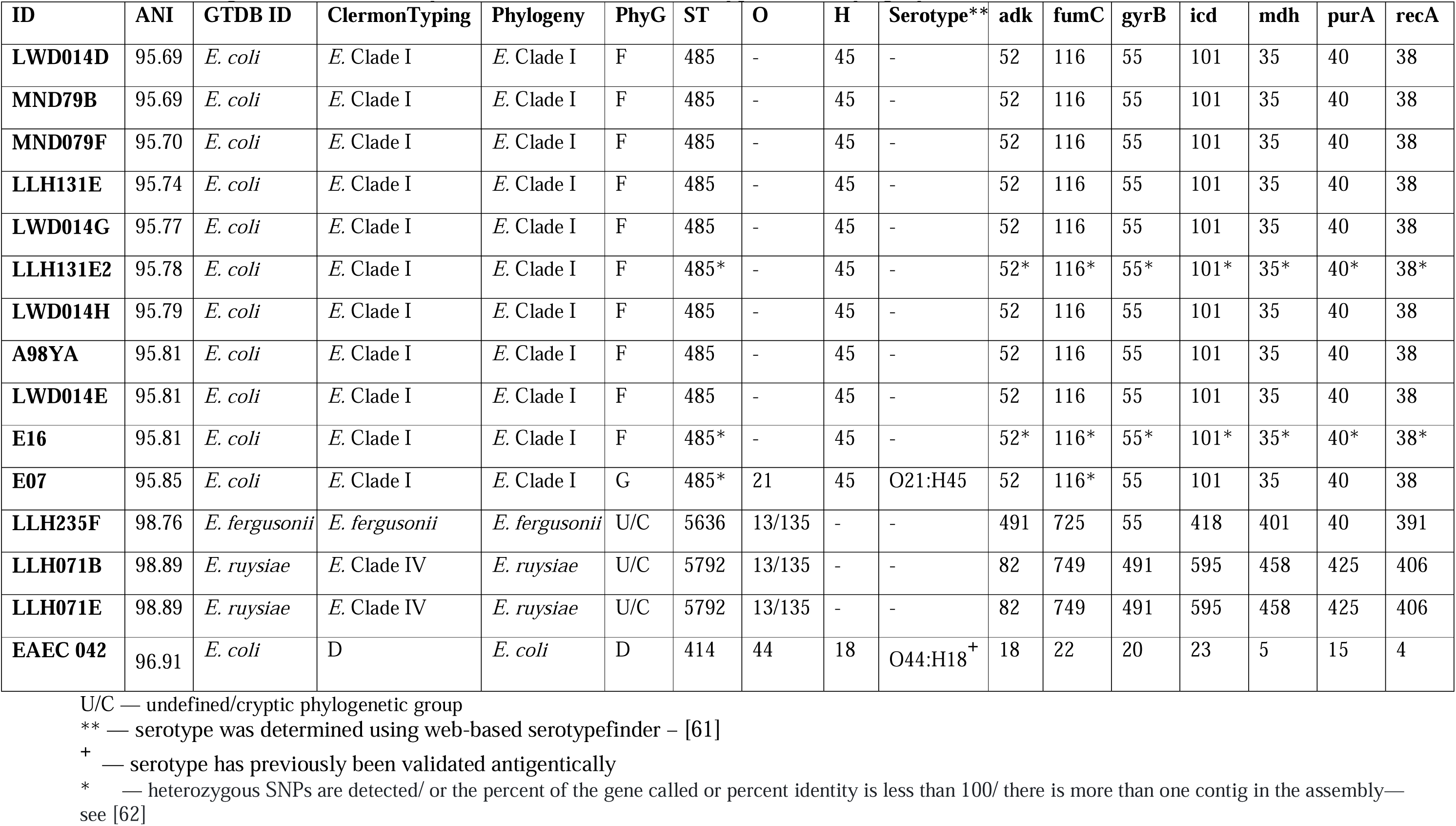
Average Nucleotide Identity (ANI) and GTDB/ClermonTyper/SNP Phylogeny ID/MLST Profiles for non-coli *Escherichia*.

When the results of GTDB-tk is compared with *E*. *coli* species abundance generated from Kraken/Bracken, contamination and completeness generated from CheckM, all the genomes had acceptable contamination and completeness values (Tab. 2). Expectedly, *E*. *fergusonii* and *E*. *ruysiae* had the fewest markers for *E. coli*, 80% and 76% respectively. As shown in Figure 1, they are phylogenetically distant from *Escherichia* Clade I. All *Escherichia* clade I genomes had between 92% and 93% *E*. *coli* genomic markers.

**Table 2:** Completeness/Contamination/Species Abundance of non-*coli Escherichia* and *Shigella* spps, *Enterobacter* spps and *S*. enterica.

| **ID** | **Completeness** | **Contamination** | ***E. coli*** | ***S. sonnei*** | ***S. boydii*** | ***S. flexneri*** | ***S. dysenteriae*** | ***Enterobacter*** | ***S. enterica*** |
| --- | --- | --- | --- | --- | --- | --- | --- | --- | --- |
| **LWD014E** | 99.7 | 0.3 | 92.2 | 0.4 | 1.8 | 0.7 | 1.1 | 1.4 | 0.6 |
| **LWD014G** | 99.7 | 0.3 | 91.5 | 0.3 | 2.0 | 0.7 | 1.2 | 1.4 | 0.6 |
| **LWD014H** | 99.7 | 0.3 | 92.7 | 0.4 | 1.1 | 0.7 | 1.0 | 1.4 | 0.6 |
| **LLH235F** | 99.9 | 0.1 | 80.2 | 0.5 | 1.8 | 0.9 | 1.2 | 4.5 | 2.3 |
| **LWD014D** | 100 | 0.2 | 91.7 | 0.4 | 1.8 | 0.7 | 1.1 | 1.3 | 0.9 |
| **MND079F** | 100 | 0.2 | 92.3 | 0.4 | 1.4 | 0.7 | 1.0 | 1.3 | 0.8 |
| **MND79B** | 100 | 0.2 | 92.3 | 0.4 | 1.4 | 0.7 | 1.0 | 1.3 | 0.8 |
| **LLH131E2** | 100 | 0.3 | 91.6 | 0.5 | 1.6 | 0.8 | 1.9 | 1.3 | 0.6 |
| **E07** | 100 | 1.1 | 92.7 | 0.4 | 1.2 | 0.8 | 1.1 | 1.5 | 0.6 |
| **E16** | 100 | 5.4 | 92.5 | 0.5 | 1.4 | 0.7 | 1.0 | 1.3 | 0.6 |
| **A98YA** | 100 | 0.2 | 92.7 | 0.3 | 1.0 | 0.7 | 1.1 | 1.4 | 0.6 |
| **LLH071B** | 100 | 0.01 | 75.9 | 1.9 | 2.7 | 1.8 | 1.8 | 5.9 | 2.0 |
| **LLH071E** | 100 | 0.01 | 76.2 | 1.8 | 2.7 | 1.8 | 1.8 | 5.8 | 2.0 |
| **LLH131E** | 100 | 0.3 | 92.1 | 0.5 | 1.6 | 0.7 | 1.0 | 1.4 | 0.5 |

One enteroaggregative *Escherichia* Clade I strain serotyped as O21:H45, using EcTyper tool. The other 13 had indeterminate O types but carried H45 flagella antigens. *E. rusiae* O types were 13/135 and their H-types were indeterminate. A subset of the *Escherichia* strains were profiled using the Microbact® system. Supplementary Table 1 shows phylogenetic groups, Microbact ID and O:H serotypes of these non-*coli Escherichia*. Of the non-coli isolates tested, Microbact identified eight as *E. coli*, one as *Hafnia alvei* and one as *Escherichia hermanii.* No test results (positive or negative) were specific or predominantly associated with non-*coli Escherichia*.

*E. coli* Clade 1 strains were between 52067 and 132638 SNPs away from the genome of *E. coli* 042 but all of them were within 700 SNPs of each other and Escherichia clade 1 reference genomes. *E. ruysiae* genomes (LLH071B and LLH071E) was, by SNP distances, the most divergent from *E. coli* reference genomes (Supplementary Table 2). Genes encoding resistance to antimicrobials of the ß-lactamase, aminoglycoside, tetracycline, sulphonamide and phenicol classes were found on the genomes of non-*coli Escherichia* in this study. The highest number of resistance genes was carried by non-*coli Escherichia* strain recovered from an adult with diarrhoea A98YA (n=20) and the fewest (n=7) by LLH131E from a healthy child and LWD014E, LWD014G, LWD014H, MND079F and MND79B, from children with diarrhoea. Figure 6 also shows the plasmid replicon types found on these genomes. Notably, *Escherichia ruysiae* genomes bore no plasmid replicon, while *p*O111 replicon was detected only in the *Escherichia fergusonii* genome. All *Escherichia* clade I genomes carried IncFII replicons. Col440I, Col8282, ColpVC were exclusively found in *Escherichia* Clade I genome E16, while rep22, rep1, rep7a and repUS1 were only carried by *Escherichia* Clade I genome A98YA.

All enteroaggregative *Escherichia* Clade I genomes contain *aap, aar* and *air*, EAEC virulence genes. They also contain virulence genes seen in other pathogenic *E. coli*, *hlyE*, *hlyF, eatA, tsh*, *traT, astA* and *chuA. Escherichia fergusonnii* and *Escherichia ruysiae* lack EAEC-specific virulence genes and had fewer virulence genes (Virulencefinder database; [63]). *Escherichia fergusonii* had only 6 virulence genes (*terC*, *gad*, *iss*, *aslA*, nlpI, and *sitA*), while *Escherichia ruysiae* had 23 virulence genes, including genes seen in *Shigella*/ enteroinvasive *E. coli* (*shiA*, *yehA*, *yehB*, *yehD*, *yghJ*, *terC, sat, chuA, fimH, iutA, csgA, AslA, anr, fdeC, cea, gad, pic, kpsE, sat, iucC, kpsMII, nlpI and traT*), but not invasion plasmid genes.

To determine whether enteroaggregative *Escherichia* clade I is found outside of our Nigeria context, we searched Enterobase [64] for ST485 genomes. As of November 14, 2025, there were 47 genome entries not associated with the current study (all labelled *E*. *coli*) uploaded between 2007 and 2025. Twenty-nine (62%) were isolated in Africa (Gambia 14; Mali 4; Kenya, 3; Madagascar 2; Mozambique 4; Tanzania 1; Zimbabwe 1), four (9%) in Europe, ten (21%) in Asia, three (6%) in the Americas, one (2%) had no information on the country of isolation. Forty-one of the enteroaggregative *Escherichia* Clade 1 strains were human isolates and one each was from a food source and a companion animal. There was no information about the source of the remaining four isolates. Some of the genomes deposited on Enterobase did not have associated accession numbers and therefore were excluded from downstream analyses. We re-interrogated all 39 ST485 genomes deposited on Enterobase which had their accession numbers provided and not associated with this study (Table 6) for the presence of virulence genes using the Virulencefinder database as reference. All 39 ST485 that we re-interrogated, regardless of their source of isolation and country/continent of isolation carried at least one EAEC-associated virulence gene (Table 6).

**Table 6:**
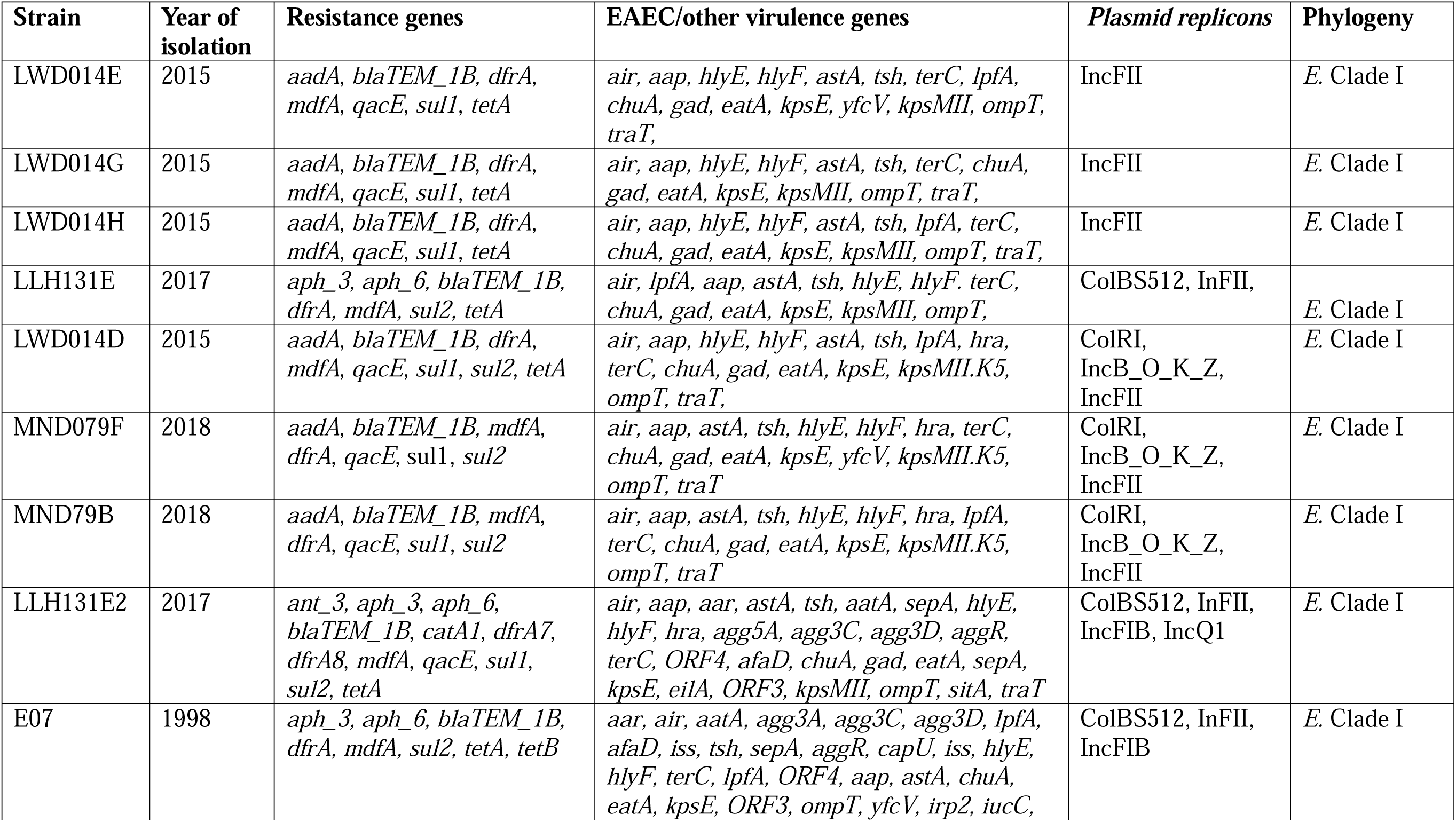
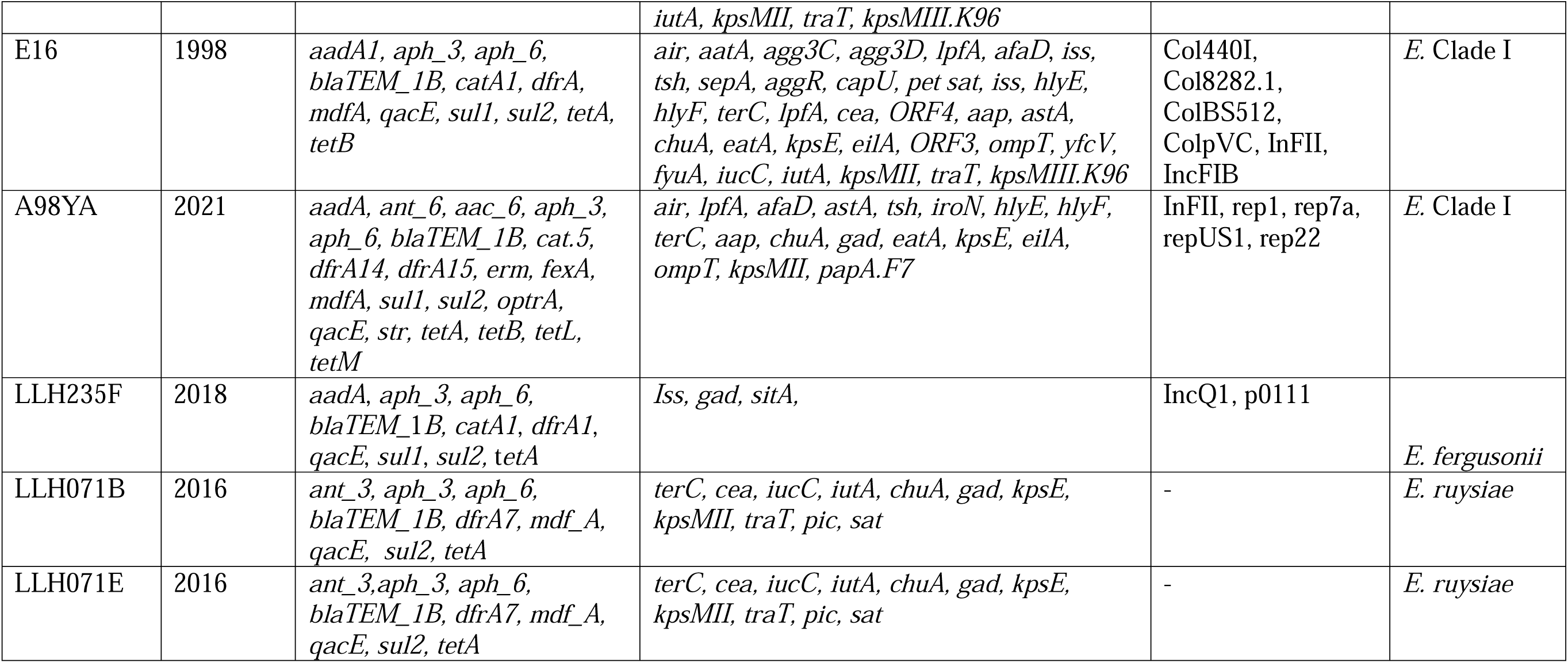
Resistance determinants, EAEC virulence genes and plasmid replicons carried by non-*coli Escherichia*.

**Table 6:** Summary of ST485 Genomes from Enterobase and their EAEC virulence genes.

| S/No | Accession Number | Source of Isolation | Continent | Country | Year of Isolation | EAEC virulence genes |
| --- | --- | --- | --- | --- | --- | --- |
| 1 | DRR387430 | Human | Asia | Japan | 2020 | *air* |
| 2 | ERR10391380 | Nil | Africa | Madagascar | Nil | *air, aap* |
| 3 | ERR10391402 | Nil | Africa | Madagascar | Nil | *air, aap* |
| 4 | ERR1790649 | Nil | Asia | Vietnam | Nil | *air* |
| 5 | ERR1790650 | Nil | Asia | Vietnam | Nil | *air* |
| 6 | ERR4554550 | Human | Europe | Netherlands | 2013 | *air* |
| 7 | ERR4557043 | Human | Europe | Netherlands | 2013 | *air, aap* |
| 8 | SRR11848721 | Human | Africa | Gambia | Nil | *air, aap, capU* |
| 9 | SRR11848781 | Human | Asia | Bangladesh | Nil | *air, aap* |
| 10 | SRR11848807 | Human | Asia | Bangladesh | Nil | *air, aap, eilA* |
| 11 | SRR11848858 | Human | Asia | India | Nil | *air, aap* |
| 12 | SRR11848878 | Human | Africa | Gambia | Nil | *air, aap, capU* |
| 13 | SRR11848919 | Human | Africa | Gambia | Nil | *air, aap, eilA, capU* |
| 14 | SRR11849099 | Human | Africa | Kenya | Nil | *air, aap, eilA* |
| 15 | SRR11849102 | Human | Africa | Gambia | Nil | *air, aap, capU* |
| 16 | SRR11849112 | Human | Africa | Mali | Nil | *air, aap* |
| 17 | SRR11849119 | Human | Africa | Mali | Nil | *air, aap* |
| 18 | SRR11849145 | Human | Africa | Mali | Nil | *air, aap, capU* |
| 19 | SRR11849161 | Human | Africa | Mozambique | Nil | *air, aap* |
| 20 | SRR11849165 | Human | Africa | Mozambique | Nil | *air* |
| 21 | SRR11849196 | Human | Africa | Mozambique | Nil | *air, aap* |
| 22 | SRR1186710 | Human | Africa | Tanzania | 2009 | *air, aap* |
| 23 | SRR12495206 | Human | Africa | Gambia | 2018 | *air, aap* |
| 24 | SRR12495207 | Human | Africa | Gambia | 2018 | *air, aap* |
| 25 | SRR12495209 | Human | Africa | Gambia | 2018 | *air, aap* |
| 26 | SRR12495210 | Human | Africa | Gambia | 2018 | *air, aap* |
| 27 | SRR21143804 | Companion animal | South America | Ecuador | 2021 | *air* |
| 28 | SRR25035178 | Food | Asia | Cambodia | 2022 | *air* |
| 29 | SRR32301530 | Human | Europe | Germany | 2016 | *air* |
| 30 | SRR32501897 | Human | Europe | United Kingdom | 2025 | *air, aatA, aaiC, aap, agg4A, aggR* |
| 31 | SRR35329232 | Human | Africa | Gambia | 2008 | *air, aap* |
| 32 | SRR35329244 | Human | Africa | Gambia | 2008 | *air, aap* |
| 33 | SRR35332962 | Human | Asia | Bangladesh | 2007 | *air, aap* |
| **34** | SRR35332963 | Human | Asia | India | 2007 | *air, aap* |
| **35** | SRR35332964 | Human | Africa | Kenya | 2008 | *air, aap* |
| **36** | SRR35332965 | Human | Africa | Mozambique | 2008 | *air, aap* |
| **37** | SRR35332982 | Human | Africa | Mali | 2008 | *air, aap* |
| **38** | SRR3588897 | Human | North America | United States | 2014 | *air, aap* |
| **39** | SRR3932419 | Human | North America | United States | 2014 | *air, aap* |

All 39 Enterobase ST485 genomes clustered with *Escherichia* clade I in our phylogeny. All but two (ERR4554550—Mash group A and ERR4557043—Mash group B1) of these genomes belong to Mash group clade I, (Figure 2). While Mash grouping use shared kmers to classify genomes, Clermontyping and Ezclermont use specific markers for classifying *Escherichia* genomes. 31 out of the 39 Enterobase ST485 genomes belong to phylogroup F (Ezclermont). Four belong to phylogroup B2, three to phylogroup G and one, to phylogroup E.

**Figure 2:**
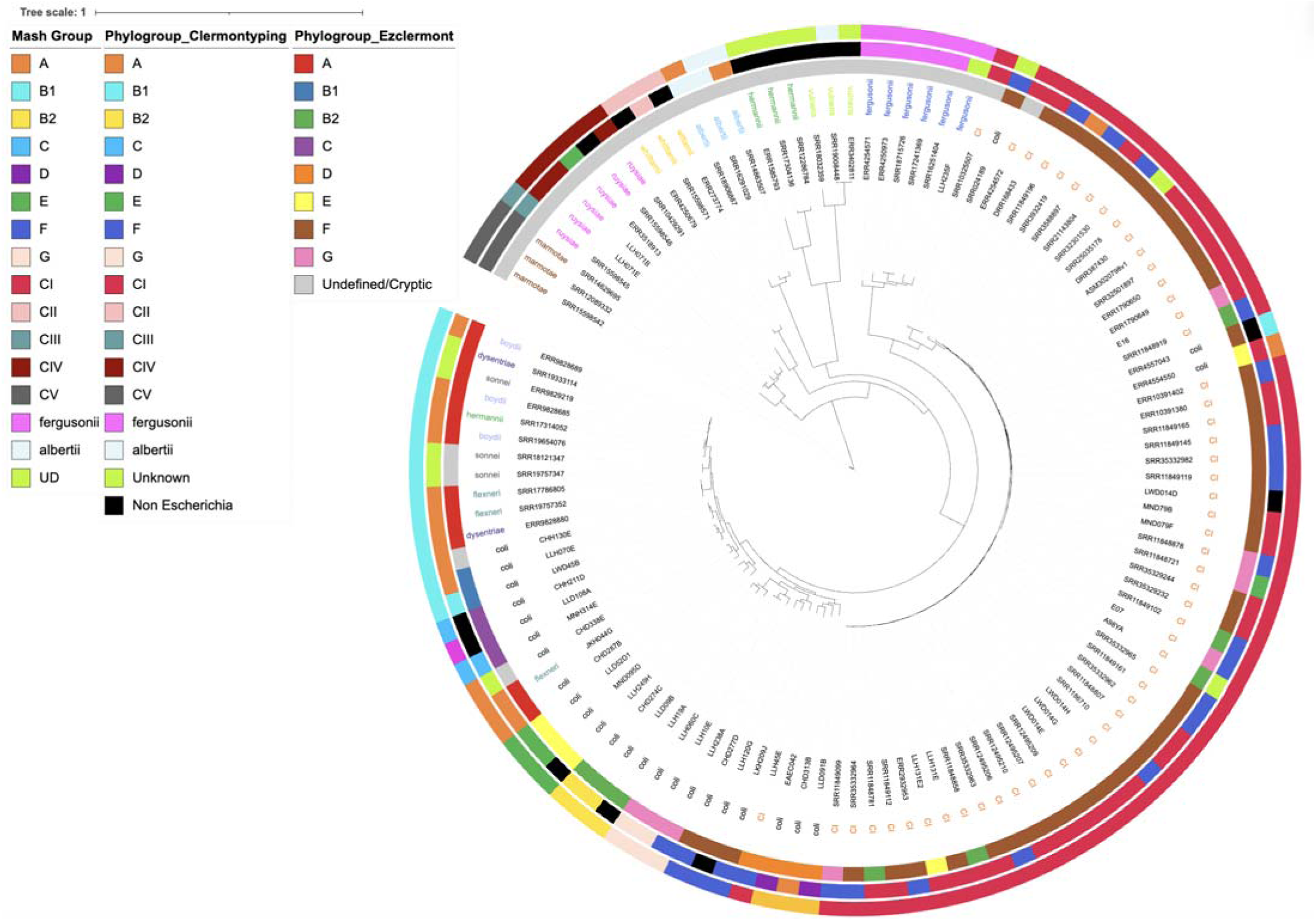
A maximum-likelihood phylogenetic tree showing mash group, clermontyping and Ezclermont phylogroups for enteroaggregative *Escherichia* ST485 from this and other studies with *E. coli* and other reference genomes

One genome (LLH45E) from our study that clustered with *E. coli* was identified as mash group clade I (ClermonTyping), while two enterobase ST485 genomes (ERR4554550 and ERR4557043) which clustered with *Escherichia* clade I (phylogeny), were identified as phylogroups A and B1 respectively. Contrarily, one genome (SRR024189) derived from NCBI’s GenBank, clustered with *Escherichia* clade I, but its mash group was undetermined (Figure 3).

**Figure 3:**
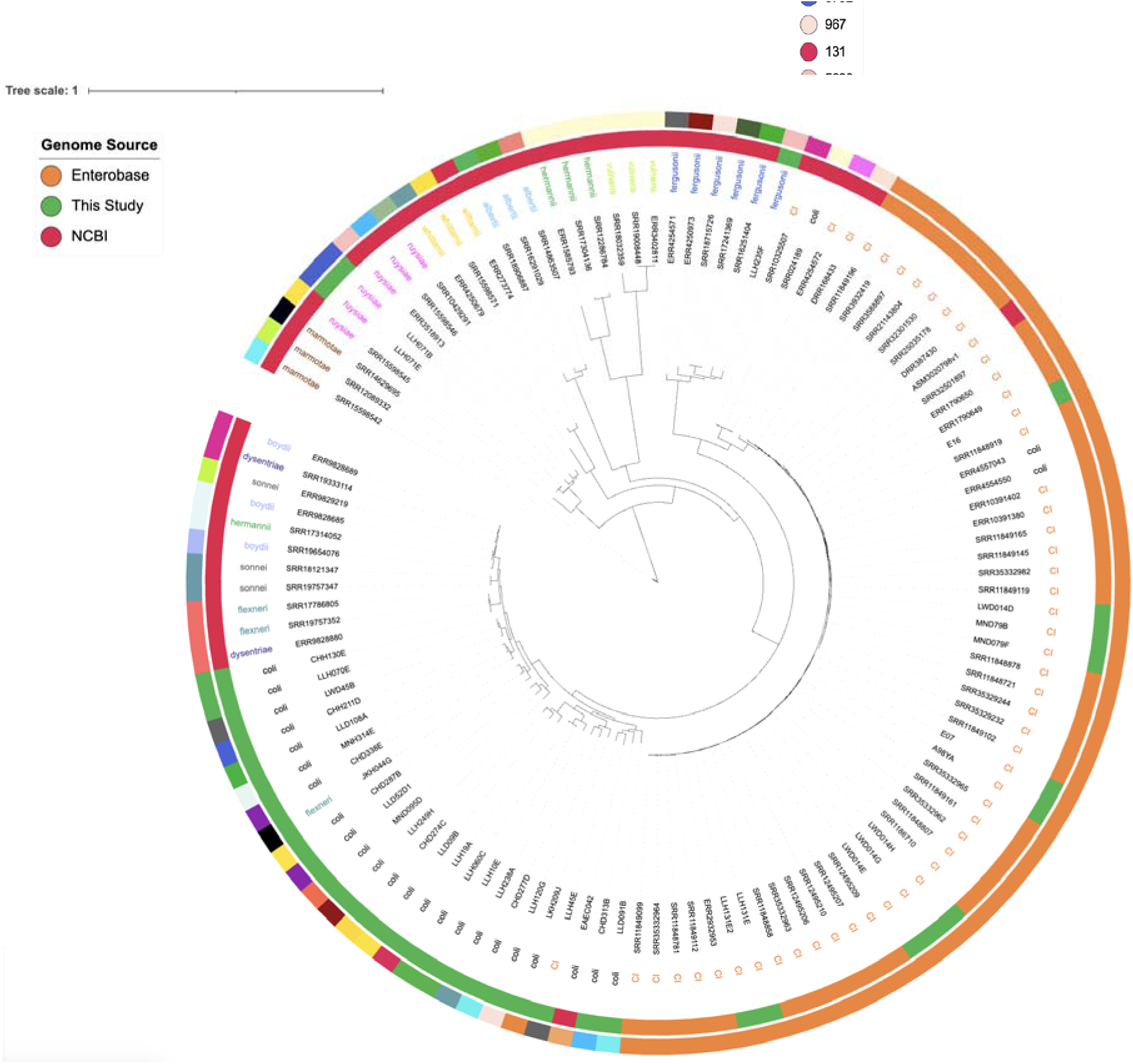
A maximum-likelihood phylogenetic tree showing ST of *E. coli* and non coli *Escherichia* from our collection, Enterobase ST485 and reference and non-*coli Escherichia*.

We also searched Enterobase [64] for ST5792 and ST5636. Genomes with this STs were identified as *E*. *ruysiae* and *E. fergusonii* in this study. Supplementary Tables 3 and 4 show the summary of genomes of ST5792 and ST5636 respectively found on Enterobase.

Figure 4 shows a goeBURST distance for MLST profiles of our non-*coli* and *coli* spp. *Escherichia*, Enterobase ST485 genomes and other reference genomes. *Escherichia fergusonii* belong to STs 5298, 5636, 7966, 10187 and 12029; *Escherichia whittamii* ST5362, ST5615 and ST11509; *Escherichia ruysiae* ST4103, ST5792, ST6467, ST9287 and ST10441; *Escherichia albertii* ST4170, ST5390 and ST12283; *Escherichia marmotae* ST133, ST5540 and ST7416; *Escherichia hermannii* ST243; *Shigella* species ST145, ST148, ST152, ST243, ST245, ST252 and ST7027; *Escherichia* Clade I ST485, ST747; ST770, ST5584 and ST8746, *Escherichia coli* STs 38, 58, 73, 90, 117, 131, 155, 206, 216, 349, 410, 423, 457, 648, 703, 967, 3018, 8318, 8489, and 15670.

**Figure 4:**
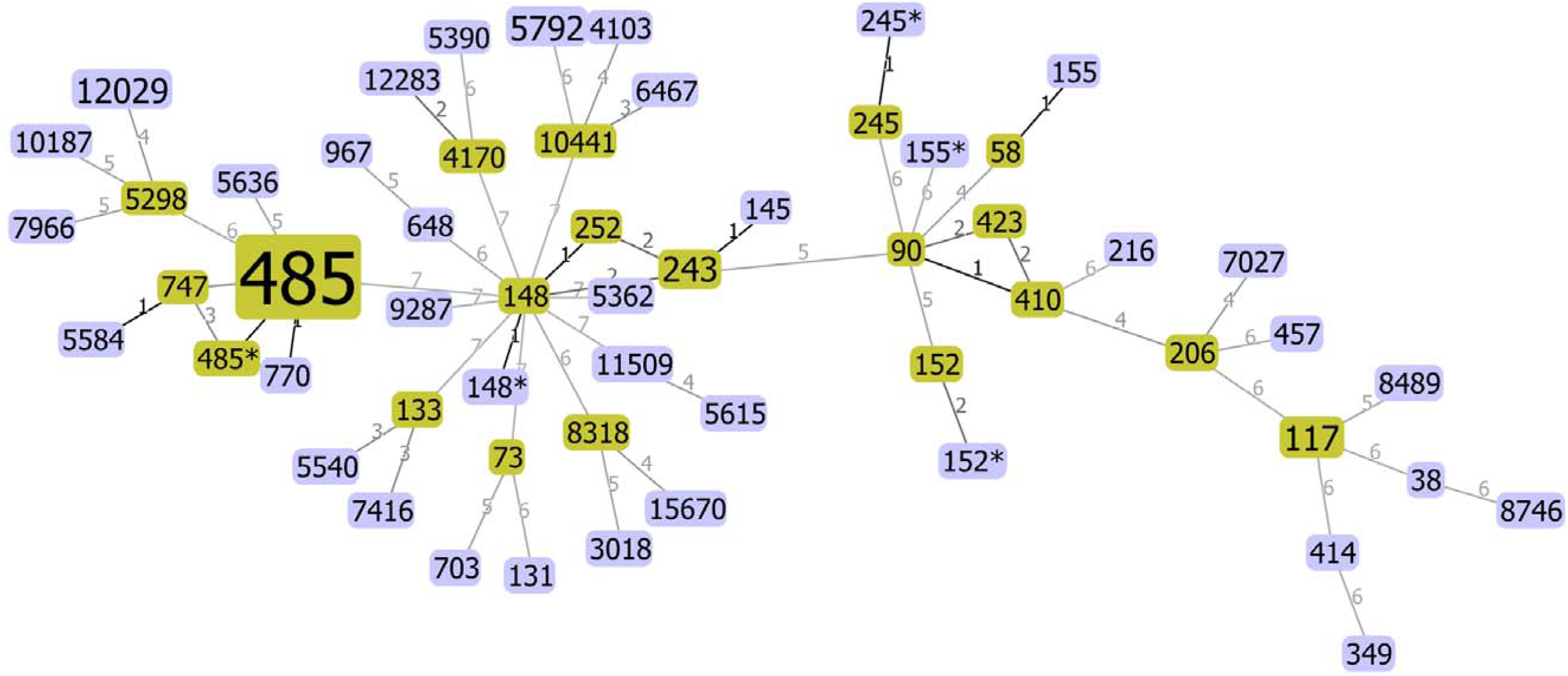
A goeburst image generated with PhyloViz software using the goeBURST algorithm (Fransisco et al., 2009) showing relatedness between STs. Each node corresponds to a different MLST profile. Founder STs are coloured green and common nodes are coloured blue. Links drawn in black connects SLVs or the same STs with ambiguity in the assembly of loci (please see notes on table 1). Dark grey links connect DLVs and light grey links connect TLVs or more. The number between lines connecting two STs represents the number of alleles by which any two connecting ST differ. Genomes with incomplete MLST alleles have been excluded.

All *Escherichia* clade I and *E*. *fergusonii* genomes regardless of their source and ST, cluster on the same branch of the Hamming distance generated by neighbour-joining using Studier-Keppler criterion. All *E. fergusonii* and *E*. clade I formed distinct sub-branches on this branch (Figure 5).

**Figure 5:**
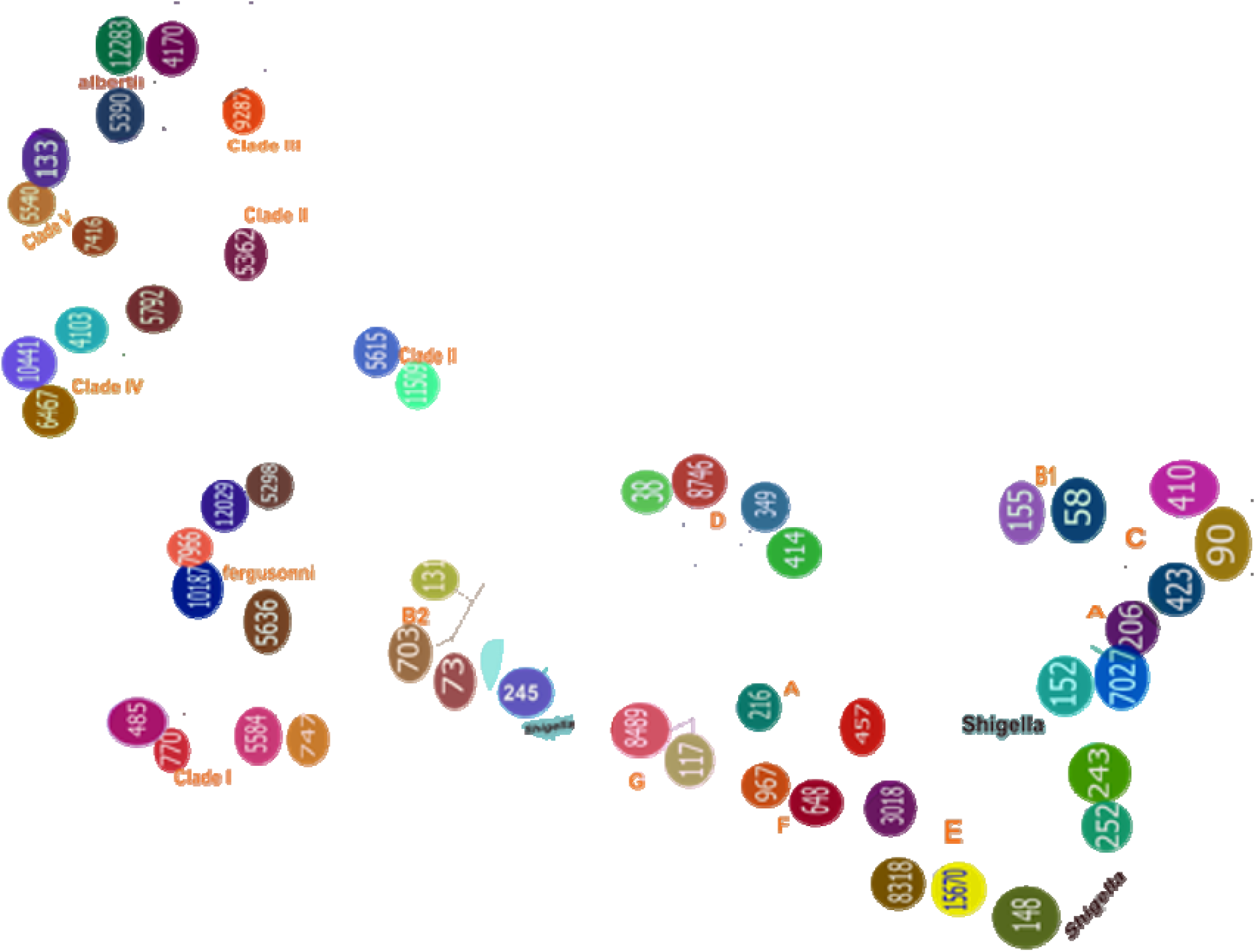
Hamming distance from MLST data of *Escherichia coli*, *Shigella* species and non *coli Escherichia* by Neighbour-Joining using Studier-Keppler criterion. ST designation with asterik (*) have been removed from this tree because of the uncertainty in the call made for the determination of their ST—[62]

## Discussion

We hypothesized that some bacteria considered as enteroaggregative *E. coli* (EAEC) belong to species other than *E. coli* and determined the species of 2,818 *Escherichia* genomes from diarrhoea epidemiology studies in Nigeria. Eleven of 1,295 putative enteroaggregative *E. coli* from epidemiology studies in Nigeria were found to be *Escherichia* Clade 1 strains and represented all but three of the non-coli isolates recovered from human stool samples. Whole Genome-based speciation analyses conducted using GTDB-tk and ClermonTyper were comparable to each other and concordant with results from the phylogeny that we generated.

Non-*coli Escherichia* species are closely related to *E. coli.* The ANI of *Escherichia* clade I isolates in this study, when compared with *E*. *col,i* showed a value of between 95.69% to 95.81%. *E*. *ruysiae* and *E*. *fergusonii* ANIs were between 98.76%. and 98.89% respectively when their nucleotides were compared with genomes within their species. Some authors have proposed the use of >96% ANI values for classifying genomes into the same species when 2 genomes are compared [65,66] and others propose higher cut-offs. [67,68,69,70].

All *Escherichia* clade I identified in this study were assigned to ST485 using the Achtman 7-loci MLST scheme and typed as *E. coli* phylogroup F or (as in one case), G using Ezclermont *E. coli* phylotyping tool. We found records of 39 other ST485 ‘*E. coli*’ in Enterobase, all closely related to enteroaggregative *Escherichia* clade I from this study. These isolates were submitted from around the world but most the genomes came from isolates recovered in Africa. This is the case even though Africa isolate genomes are under-represented in Enterobase, with only 9,971 of 316,247 *E. coli* genomes originating from Africa as of 10 December 2025).

This study additionally identified three other non-coli *Escherichia* from the Nigeria strain set and from Enterobase at lower frequencies. We identified two ST5792 as *E*. *ruysiae* and there were 11 ST5792 genomes on Enterobase. Four of these from human sources, one from the environment and the remaining six had no information on their source of isolation. *E. ruysiae* human infections are rarely reported and there are few reports linking *E*. *ruysiae* to human or animal infections [16,22,71,72]. We also found one ST5636 genome in this study which we speciated as *E. fergusonii* using SNP phylogeny. Our Enterobase search revealed that there were 21 ST5636 submissions as of December 10, 2025. All but one genome (submitted by the Canadian Food Inspection Agency from food source and indexed as *E. fergusonii*) had no information on their source of isolation, accession number and species designation. *E. fergusonii* has been reported from blood, urine and faeces [17,20].

Zhang and his colleagues carried out a comparative genetic characterisation of EAEC strains recovered from both clinical (51 isolates) and non-clinical (33 isolates) settings. Using PCR and the 7-loci housekeeping genes, they recovered one ST485 EAEC genome from a child with diarrhoea [73]. In a study carried out by Foster-Nyarko and his colleagues to determine the genomic diversity of *E*. *coli* isolated from healthy children in the Gambia, one (out of 88) ST485 *Escherichia* clade I isolate was recovered [74]. Their ST485 carried among other virulence genes, *air* and *aap* which appears to be typically present in most *Escherichia* clade I genomes. Using PCR, Zueter and colleagues carried out a multilocus sequence typing of *E. coli* isolated from clinical samples in Jordan and found one ST485 in their study [75]. They did not determine the virulence-associated genes carried by their genomes making it impossible to determine if their ST 485 genome carried any enteroaggregative genes and if they belong to *Escherichia* clade I. We however can make inference that their ST485 might be *Escherichia* clade I owing to its position on their phylogenetic tree which was constructed using concatenated sequences of every allelic profile, joined in the order of the alleles that they used for the determination of MLST.

While we have shown that all our *Escherichia* clade I genomes belong to ST485 and that ST485 potentially might be the most predominant clade I ST, there may be other enteroaggregative *Escherichia* clade I STs. For example, a study carried out by Hoshiko and others on 91 *E. coli* genomes associated with human diarrhoea from India, one strain was classified as *Escherichia* clade I. This strain belongs to ST5584 and serotype O17:H45, sharing the same flagellar antigen type with *Escherichia* clade I found in this study. This strain also carried among other virulence genes *eilA* which is associated with EAEC and *elt1*/*estA* which are used to define ETEC. Additionally, the clade I strain from their study also carried plasmid replicons (Col440I and IncFIB) found in some of the *Escherichia* clade I in this study [76]. Aworh and her colleagues [77], to determine the genetic relatedness of multidrug resistant *E*. *coli* isolated from humans, chickens and poultry environments in Abuja, identified one *Escherichia* clade I, ST5584 isolate and one *Escherichia* clade IV (*E*. *ruysiae*), ST10441 isolate. Both non-*coli Escherichia* were recovered from chicken farms. Their *Escherichia* clade I isolate carried genes *air* and *eilA* which are associated with EAEC [49,78] and gene *hra* which is an accessory colonisation factor in prototypical EAEC 042 [79]. Additionally, the clade I that they isolated differs from ST485 clade I by four MLST loci, sharing only *adk*, *recA* and *purA* loci with ST485 [77]. *Escherichia* clade I belonging to other STs other than ST485 have also been reported elsewhere. Tiwari and colleagues carried out a Genome-wide association study to understand host-specific genomic traits in *Escherichia coli* over a 16-year period. Their genomes were derived from isolates from Vietnam, UK, Germany and Spain and from a variety of hosts (wild boar, chicken, human, cattle and pig). Of the 1198 whole genome sequenced isolates from healthy and diseased hosts, eight *Escherichia* clade I belonging to ST770 (n=6) from chickens, ST3057 (n=1) from pig and ST5273 (n=1) from chicken were found—all from healthy hosts. In their study all *Escherichia* clade I were of non-human origin, were not involved in disease and do not belong to ST485 [80]. ST770 is a single locus variant of ST485, differing at the mdh MLST locus. ST3057 and ST485 share 3 loci, while ST5273 shared no loci with ST485, but have 2 common loci with ST3057. Interestingly, all the eight clade I genomes that they recovered from their isolates carried genes *air* and *eilA* which are EAEC-associated. This may indicate that *Escherichia* clade I, regardless of their MLST assignment are non-*coli* enteroaggregatives.

Wang and others carried out a study to understand the prevalence and genetic characteristics of pathogenic *E. coli* in pigeon farms, in central China. They identified 11 sequence types and one out of 60 *E*. *coli* that they isolated belong to ST485 [81].While they classified this isolate into phylogenetic group B2, it had H-antigen 45 (H45), which we found on all our *Escherichia* clade I genomes. Interestingly, all non clade I *Escherichia* in this study had untypable O antigens. Wang and his co-workers determined the virulence factors carried by their genomes, but only genes that regulate iron uptake were reported. Other virulence genes, including those that define EAEC were not reported.Yang and his colleagues also carried out a study to determine the frequency of virulence genes and characteristics of diarrheagenic in *E. coli* isolated from pigs with diarrhoea in China and reported that they found one ST485 among their collection of 65 STs from 171 isolates [82]. They used real-time PCR assays for the detection of 11 virulence genes, but none of the virulence genes that they targeted are EAEC-associated. However, their ST485 carried genes that code for LT/STb and Stx2e. These toxins are important virulence factors for enterotoxigenic *E*. *coli* and enterohaemorrhagic *E*. *coli* respectively [42,83]. When we interrogated their genome (ASM3020798v1), we found that it belongs to clade I and carried genes *air* (an enteroaggregative gene) and *hlyE* which is associated with APEC.

We attempted to see if we could use Microbact Octal number and percentage probability results generated to define a threshold for identifying suspected *Escherichia* clade I. We did not find any pattern that might be employed to presumptively identify suspected *Escherichia* clade I in routine laboratories where there is no capacity for molecular testing or genome sequencing and bioinformatics. Microbact 24E is able to identify “*E*. *coli* inactive” but strains returned as “*E*. *coli* inactive” were identified as *E*. *coli* using whole genome SNPs phylogeny and were spread across various phylogenetic groups including phyloroups A, B2, C and F.

We observed that *Escherichia* clade I genomes from this study are more similar to one another, based on SNP differences, than are *E. coli* lineages. This was the case even for two isolates (E07, E18) that were isolated over two decades ago [23], in a different Nigerian state, as well as for *Escherichia* clade 1 human isolates in the database. Curiously, *Escherichia* clade I appears to be a stable diarrhoeagenic clone over nearly 30 years, being associated with diarrhoea in children then, and more recently.

The possession EAEC-specific factors including the gene *air* that codes of enteroaggregative immunoglobulin repeat protein, gene *aap* that codes for anti-aggregation protein (dispersin) and other EAEC-associated virulence genes clearly distinguishes the *Escherichia* clade I from other non-*coli Escherichia* which lack any of these EAEC genes. Of note is our observation that the presence of EAEC-associated virulence factors in *Escherichia* clade I (regardless of their MLST profile, niche or geographical location) and the absence of these EAEC defining genes in other non-*coli Escherichia*. This observation was also true for all *Escherichia* clade I and other non-*coli Escherichia* retrieved from Enterobase.

In our experience, SNP phylogeny is a reliable tool for defining *Escherichia* clade I from whole genome sequence data. And of the duo, (Ezclermont and Clermontyping), Clermontyping provides a comparator which uses Mash to determine the species of *Escherichia* clade I. This provides two layers of results (phylogenetic group and mash group), which can be used for resolving results when there is disconcordance in the final ouput generated by Clermontyping and Ezclermont. This supplementation of phylogrouping with Mash grouping in Clermontyping provided speciation results that agreed with the results from SNP phylogeny for *Escherichia* clade I.

Our MLST data and the resulting analyses carried out using the goeBURST algorithm on PhyloViz confirmed the homogeneity of *Escherichia* clade I, their niches notwithstanding. We also showed (from our data and the data that we retrieved from other studies) that Escherichia clade I may have evolved from *E*. *fergusonii*: ST5636 *Escherichia* clade I and formed a distinct sub-branch on the cladeI-*fergusonii* branch of the Hamming distance generated from MLST data. This relatedness was also confirmed with SNPs phylogeny using SNPs from whole genome sequenced data.

All 11 *Escherichia* clade I in this study were assigned to ST485. Our findings from our own data and from the Enterobase database suggest that ST485 might be the predominant ST among *Escherichia* clade derived from human sources and that other clade I STs may be more associated with other niches. That Escherichia Clade I ST485 strains are largely overlooked, even by enteroaggregative *E. coli* researchers may reflect the neglected status of this pathotype, and their predominance on the African continent, from which few diarrhoeagenic *Escherichia* are identified or sequenced.

## Acknowledgements

We thank El-shama Q. Nwoko, Stephen O. Bejide, Pelumi Daniel Adewole, Jola-Ade Ajiboye and David A Kwasi for technical and administrative assistance.

## List of Supplementary Materials

1. **Supplementary Table 1:** Phylogenetic group and Microbact ID of *coli* and non-*coli Escherichia*
2. **Supplementary Table 2:** Core SNP differences between putative non-*coli Escherichia* and reference genomes
3. **Supplementary Table 3**: Summary of ST5792 Genomes found on Enterobase
4. **Supplementary Table 4:** Summary of ST5636 Genomes found on Enterobase

## References

1. Koh XP, Shen Z, Woo CF, Yu Y, Lun HI, Cheung SW, et al. Genetic and Ecological Diversity of *Escherichia coli* and Cryptic *Escherichia* Clades in Subtropical Aquatic Environments. Front. Microbiol. 2022;13:811755. 10.3389/fmicb.2022.811755

2. Pokharel P, Dhakal S, Dozois CM. The Diversity of *Escherichia coli* Pathotypes and Vaccination Strategies against This Versatile Bacterial Pathogen. Microorganisms. 2023;11(2):344. 10.3390/microorganisms11020344

3. Thomson NM, Gilroy R, Getino M, Foster-Nyarko E, van Vliet AH, La Ragione RM, et al. Remarkable genomic diversity among *Escherichia* isolates recovered from healthy chickens. PeerJ. 2022;10:e12935 DOI 10.7717/peerj.12935

4. Yu D, Banting G, Neumann NF. (2021). A review of the taxonomy, genetics, and biology of the genus *Escherichia* and the type species *Escherichia coli*. Can J. Microbiol. 2021;67(8):553-571. 10.1139/cjm-2020-0508

5. Lagerstrom KM, Hadly EA. Under-Appreciated Phylogroup Diversity of *Escherichia coli* within and between Animals at the Urban-Wildland Interface. Appl Environ Microbiol. 2023;89(6):e0014223. doi:10.1128/aem.00142-23

6. Braz VS, Melchior K, Moreira CG. *Escherichia coli* as a Multifaceted Pathogenic and Versatile Bacterium. Front. Cell. Infect. Microbiol. 2020;10:548492. 10.3389/fcimb.2020.548492

7. Koh XP, Shen Z, Woo CF, Yu Y, Lun HI, Cheung SW, et al. Genetic and Ecological Diversity of *Escherichia coli* and Cryptic *Escherichia* Clades in Subtropical Aquatic Environments. Front Microbiol. 2022;13:811755. doi: 10.3389/fmicb.2022.811755.

8. Okuno M, Tsuru N, Yoshino S, Gotoh Y, Yamamoto T, Hayashi T, et al. Isolation and Genomic Characterization of a Heat-Labile Enterotoxin 1-Producing *Escherichia fergusonii* Strain from a Human. Microbiol Spectr. 2023;11(4):e0049123. doi: 10.1128/spectrum.00491-23.

9. Walk ST. The “Cryptic” *Escherichia*. EcoSal Plus. 2015;6(2), 10.1128/ecosalplus.ESP-0002-2015. 10.1128/ecosalplus.ESP-0002-2015

10. Ioannou P. *Escherichia hermannii* Infections in Humans: A Systematic Review. Trop Med Infect Dis. 2019;4(1):17. doi: 10.3390/tropicalmed4010017.

11. Halimeh FB, Rafei R, Osman M, Kassem II, Diene SM, Dabboussi F, et al. Historical, current, and emerging tools for identification and serotyping of *Shigella*. Braz. J. Microbiol. 2021;52(4), 2043–2055. 10.1007/s42770-021-00573-5

12. Cobo-Simón M, Hart R, Ochman H. *Escherichia coli*: What Is and Which Are?. Mol Biol Evol. 2023;40(1):msac273. doi:10.1093/molbev/msac273

13. Lan R, Reeves PR. Escherichia coli in disguise: molecular origins of *Shigella*. Microbes Infect. 2002;4(11):1125–1132. doi:10.1016/s1286-4579(02)01637-4

14. Chattaway MA, Schaefer U, Tewolde R, Dallman TJ, Jenkins C. Identification of *Escherichia coli* and *Shigella* Species from Whole-Genome Sequences. J. Clin. Microbiol 2017;55(2): 616–623. 10.1128/JCM.01790-16

15. Sinha T, Merlino J, Rizzo S, Gatley A, Siarakas S, Gray T. Unrecognised: isolation of *Escherichia marmotae* from clinical urine sample, phenotypically similar to *Escherichia coli*. Pathology. 2024;56(4), 577–578. 10.1016/j.pathol.2023.08.015

16. van der Putten BCL, Matamoros S, Mende DR, Scholl ER, Consortium C, Schultsz C. *Escherichia ruysiae* sp. nov., a novel Gram-stain-negative bacterium, isolated from a faecal sample of an international traveller. Int J Syst. Evol. Microbiol. 2021;71(2):004609. doi: 10.1099/ijsem.0.004609.

17. Savini V, Catavitello C, Talia M, Manna A, Pompetti F, Favaro M, et al. Multidrug-Resistant *Escherichia fergusonii*: a Case of Acute Cystitis. J. Clin. Microbiol. 2008;46(4):1551–1552 0095-1137/08/$08.00 0 doi:10.1128/JCM.01210-07

18. Zang YM, Liu JF, Li G, Zhao M, Yin GM, Zhang ZP, et al. The first case of *Escherichia fergusonii* with biofilm in China and literature review. BMC Infect Dis 2023;23:35. 10.1186/s12879-023-07985-8

19. Muchaamba F, Barmettler K, Treier A, Houf K, Stephan R. Microbiology and Epidemiology of *Escherichia albertii*—An Emerging Elusive Foodborne Pathogen. Microorganisms. 2022;10(5):875. 10.3390/microorganisms10050875

20. Parin U, Kirkan S, Arslan SS, Yuksel HT. Molecular identification and antimicrobial resistance of *Escherichia fergusonii* and *Escherichia coli* from dairy cattle with diarrhoea. Veterinarni Medicina, 2018;63(03):110–116 10.17221/156/2017-VETMED

21. An F, Wang K, Wei S, Yan H, Xu X, Xu J, et al. First case report of pustules associated with *Escherichia fergusonii* in the Chinese pangolin (*Manis pentadactyla aurita*). BMC Vet Res. 2023;19(1):69. doi: 10.1186/s12917-023-03622-3.

22. Dione N, Mlaga KD, Liang S, Jospin G, Marfori Z, Alvarado N, et al. Comparative genomic and phenotypic description of *Escherichia ruysiae*: a newly identified member of the gut microbiome of the domestic dog. Front Microbiol. 2025;16:1558802. doi: 10.3389/fmicb.2025.1558802.

23. Okeke IN, Lamikanra A, Steinrück H, Kaper JB. Characterization of *Escherichia* coliStrains from Cases of Childhood Diarrhea in Provincial Southwestern Nigeria. J Clin Microbiol. 2000;38. 10.1128/jcm.38.1.7-12.2000

24. Rabiu AG, Falodun OI, Fagade OE, Dada RA, Okeke IN. Potentially pathogenic *Escherichia coli* from household water in peri-urban Ibadan, Nigeria. J Water Health. 2022;20(7):1137–1149. doi:10.2166/wh.2022.117

25. Rabiu AG, Falodun OI, Dada RA, Afolayan AO, Akinlabi OC, Akande ET, Okeke IN. Transmissible antimicrobial resistance in *Escherichia coli* isolated from household drinking water in Ibadan, Nigeria. PLoS One. 2025;20(5):e0318969. doi: 10.1371/journal.pone.0318969.

26. Akinlabi OC, Dada RA, Nwoko EQA, Okeke IN. PCR diagnostics are insufficient for the detection of Diarrhoeagenic *Escherichia coli* in Ibadan, Nigeria. PLOS Glob. Public Health. 2023;3(8):e0001539. 10.1371/journal.pgph.0001539

27. Bejide OS, Odebode MA, Ogunbosi BO, Adekanmbi O, Akande KO, Ilori T, et al. Diarrhoeal pathogens in the stools of children living with HIV in Ibadan, Nigeria. Front. Cell. Infect. Microbiol. 2023;13. 10.3389/fcimb.2023.1108923

28. Olayinka AA, Oginni-Falajiki I O, Okeke IN, Aboderin AO. Diarrhoeagenic *Escherichia coli* associated with childhood diarrhoea in Osun state, Nigeria. BMC Infect. Dis. 2024;24(1):928. 10.1186/s12879-024-09793-0

29. Adewole et al (unpublished)

30. www.bioinformatics.babraham.ac.uk/projects/

31. Ewels P, Magnusson M, Lundin S, Käller M. MultiQC: summarize analysis results for multiple tools and samples in a single report. Bioinformatics. 2016;32(19), 3047–3048. 10.1093/bioinformatics/btw354

32. Wood DE, Lu J, Langmead B. Improved metagenomic analysis with Kraken 2. Genome Biol. 2019;20:257. 10.1186/s13059-019-1891-0

33. Lu J, Breitwieser FP, Thielen P, Salzberg SL. Bracken: estimating species abundance in metagenomics data. PeerJ Comput Sci. 2017;3:e104. doi:10.7717/peerj-cs.104

34. 34. https://github.com/sanger-pathogens/sh16_scripts

35. Page AJ, Cummins CA, Hunt M, Wong VK, Reuter S, Holden MT, et al. Roary: rapid large-scale prokaryote pan genome analysis. Bioinformatics. 2015;31(22):3691–3693. 10.1093/bioinformatics/btv421

36. Minh BQ, Schmidt HA, Chernomor O, Schrempf D, Woodhams MD, von Haeseler A, et al. IQ-TREE 2: New Models and Efficient Methods for Phylogenetic Inference in the Genomic Era. Mol. Biol. and Evol. 2020;37(5):1530–1534. 10.1093/molbev/msaa015

37. (https://tree.bio.ed.ac.uk/software/figtree/

38. Hadfield J, Croucher NJ, Goater RJ, Abudahab K, Aanensen DM, Harris SR. Phandango: an interactive viewer for bacterial population genomics. Bioinformatics. 2018;34(2):292–293. doi: 10.1093/bioinformatics/btx610.

39. Letunic I, Bork P. Interactive Tree Of Life (iTOL) v5: an online tool for phylogenetic tree display and annotation. Nucleic Acids Res. 2021;49(W1):W293–W296. doi: 10.1093/nar/gkab301.

40. Argimón S, Abudahab K, Goater RJE, Fedosejev A, Bhai J, Glasner C, et al. Microreact: visualizing and sharing data for genomic epidemiology and phylogeography. Microb Genom. 2016;2(11):e000093. doi: 10.1099/mgen.0.000093.

41. Guindon S, Dufayard JF, Lefort V, Anisimova M, Hordijk W, Gascuel O. New algorithms and methods to estimate maximum-likelihood phylogenies: assessing the performance of PhyML 3.0. Syst Biol. 2010;59(3):307–21. doi: 10.1093/sysbio/syq010.

42. Boisen N, Østerlund MT, Joensen KG, Santiago AE, Mandomando I, Cravioto A, et al. Redefining enteroaggregative Escherichia coli (EAEC): Genomic characterization of epidemiological EAEC strains. PLoS Negl Trop Dis. 2020;14(9):e0008613. doi: 10.1371/journal.pntd.0008613.

43. Croxen MA, Law RJ, Scholz R, Keeney KM, Wlodarska M, Finlay BB. Recent advances in understanding enteric pathogenic Escherichia coli. Clin Microbiol Rev. 2013;26(4):822–80. doi: 10.1128/CMR.00022-13.

44. Robins-Browne RM, Holt KE, Ingle DJ, Hocking DM, Yang J, Tauschek M. Are *Escherichia coli* Pathotypes Still Relevant in the Era of Whole-Genome Sequencing? Front Cell Infect Microbiol. 2016;6:141. doi: 10.3389/fcimb.2016.00141.

45. Lima IFN, Boisen N, Silva JDQ, Havt A, de Carvalho EB, Soares AM, et al. Prevalence of enteroaggregative *Escherichia coli* and its virulence-related genes in a case-control study among children from north-eastern Brazil. J Med Microbiol. 2013;62(Pt 5):683–693. doi: 10.1099/jmm.0.054262-0.

46. Sheikh J, Czeczulin JR, Harrington S, Hicks S, Henderson IR, Le Bouguénec C, et al. A novel dispersin protein in enteroaggregative Escherichia coli. J Clin Invest. 2002;110(9):1329–37. doi: 10.1172/JCI16172.

47. Czeczulin JR, Whittam TS, Henderson IR, Navarro-Garcia F, Nataro JP. Phylogenetic Analysis of Enteroaggregative and Diffusely Adherent *Escherichia coli*. Infect Immun 1999;67:.10.1128/iai.67.6.2692-2699.1999

48. Mickey AS, Nataro JP. Dual function of Aar, a member of the new AraC negative regulator family, in *Escherichia coli* gene expression. Infect Immun 2020;88:e00100–20. 10.1128/IAI.00100-20.

49. Sheikh J, Dudley EG, Sui B, Tamboura B, Suleman A, Nataro JP. EilA, a HilA-like regulator in enteroaggregative *Escherichia coli*. Mol Microbiol. 2006;61(2):338–50. doi: 10.1111/j.1365-2958.2006.05234.x.

50. Hunt M, Mather AE, Sánchez-Busó L, Page AJ, Parkhill J, Keane JA, et al. ARIBA: rapid antimicrobial resistance genotyping directly from sequencing reads. Microb Genom. 2017;3(10):e000131. doi: 10.1099/mgen.0.000131.

51. 51. https://pubmlst.org/

52. Malberg Tetzschner AM, Johnson JR, Johnston BD, Lund O, Scheutz F. *In Silico* Genotyping of *Escherichia coli* Isolates for Extraintestinal Virulence Genes by Use of Whole-Genome Sequencing Data. J Clin Microbiol. 2020;58(10):e01269–20. doi: 10.1128/JCM.01269-20.

53. Florensa AF, Kaas RS, Clausen PTLC, Aytan-Aktug D, Aarestrup FM. ResFinder - an open online resource for identification of antimicrobial resistance genes in next-generation sequencing data and prediction of phenotypes from genotypes. Microb Genom. 2022;8(1):000748. doi: 10.1099/mgen.0.000748.

54. Carattoli A, Hasman H. PlasmidFinder and In Silico pMLST: Identification and Typing of Plasmid Replicons in Whole-Genome Sequencing (WGS). Methods Mol. Biol. 2020;2075, 285–294. 10.1007/978-1-4939-9877-7_20

55. Bankevich A, Nurk S, Antipov D, Gurevich AA, Dvorkin M, Kulikov AS, et al. SPAdes: a new genome assembly algorithm and its applications to single-cell sequencing. J Comput Biol. 2012;19(5):455–77. doi: 10.1089/cmb.2012.0021.

56. Parks DH, Imelfort M, Skennerton CT, Hugenholtz P, Tyson GW. CheckM: assessing the quality of microbial genomes recovered from isolates, single cells, and metagenomes. Genome Res. 2015;25(7):1043–55. doi: 10.1101/gr.186072.114.

57. Bessonov K, Laing C, Robertson J, Yong I, Ziebell K, Gannon VPJ, et al. ECTyper: *in silico Escherichia coli* serotype and species prediction from raw and assembled whole-genome sequence data. Microb Genom. 2021;7(12):000728. doi: 10.1099/mgen.0.000728.

58. Waters NR, Abram F, Brennan F, Holmes A, Pritchard L. Easy phylotyping of *Escherichia coli via* the EzClermont web app and command-line tool. Access Microbiol. 2020;2(9):acmi000143. doi:10.1099/acmi.0.000143

59. Chaumeil P-A, Mussig AJ, Hugenholtz P, Parks DH. GTDB-Tk v2: memory friendly classification with the genome taxonomy database, Bioinformatics. 2022;38(23):5315–5316, 10.1093/bioinformatics/btac672

60. Beghain J, Bridier-Nahmias A, Le Nagard H, Denamur E, Clermont O. ClermonTyping: an easy-to-use and accurate in silico method for Escherichia genus strain phylotyping. Microb Genom. 2018;4(7):e000192. doi: 10.1099/mgen.0.000192.

61. https://cge.food.dtu.dk/services/SerotypeFinder/

62. https://github.com/sanger-pathogens/ariba/wiki/MLST-calling-with-ARIBA

63. https://bitbucket.org/genomicepidemiology/virulencefinder_db/src/master/

64. www.enterobase.warwick.ac.uk

65. Ciufo S, Kannan S, Sharma S, Badretdin A, Clark K, Turner S, et al. (2018). Using average nucleotide identity to improve taxonomic assignments in prokaryotic genomes at the NCBI. Int J Syst Evol Microbiol. 2018;68(7):2386–2392. doi: 10.1099/ijsem.0.002809.

66. Pearce ME, Langridge GC, Lauer AC, Grant K, Maiden MCJ, Chattaway MA. An evaluation of the species and subspecies of the genus Salmonella with whole genome sequence data: Proposal of type strains and epithets for novel S. enterica subspecies VII, VIII, IX, X and XI. Genomics. 2021;113(5):3152–3162. doi: 10.1016/j.ygeno.2021.07.003.

67. Palmer M, Steenkamp ET, Blom J, Hedlund BP, Venter SN. All ANIs are not created equal: implications for prokaryotic species boundaries and integration of ANIs into polyphasic taxonomy. Int J Syst Evol Microbiol. 2020;70(4):2937–2948. doi: 10.1099/ijsem.0.004124.

68. Rodriguez-R LM, Jain C, Conrad RE, Aluru S, Konstantinidis KT. Reply to: “Re-evaluating the evidence for a universal genetic boundary among microbial species”. Nat Commun. 2021;12:4060 10.1038/s41467-021-24129-1

69. Park MJ, Kim YJ, Park M, Yu J, Namirimu T, Roh YR, et al. Establishment of Genome Based Criteria for Classification of the Family Desulfovibrionaceae and Proposal of Two Novel Genera, Alkalidesulfovibrio gen. nov. and Salidesulfovibrio gen. nov. Front Microbiol. 2022;13:738205. doi: 10.3389/fmicb.2022.738205.

70. Briand M, Bouzid M, Hunaul, G, Legeay M, Fischer-Le Saux M, Barret MA. Rapid and simple method for assessing and representing genome sequence relatedness. (2021). Peer Community J. 2021;1: e24. 10.24072/pcjournal.37

71. Campos-Madueno EI, Aldeia C, Sendi P, Endimiani A. *Escherichia ruysiae* May Serve as a Reservoir of Antibiotic Resistance Genes across Multiple Settings and Regions. Microbiol Spectr. 2023;11:e01753–23. 10.1128/spectrum.01753-23

72. Siddi G, Piras F, Gymoese P, Torpdahl M, Meloni MP, Cuccu M, et al. Pathogenic profile and antimicrobial resistance of *Escherichia coli*, *Escherichia marmotae* and *Escherichia ruysiae* detected from hunted wild boars in Sardinia (Italy). Int J Food Microbiol. 2024;421:110790. doi: 10.1016/j.ijfoodmicro.2024.110790.

73. Zhang R, Gu DX, Huang YL, Chan EW, Chen GX, Chen S. Comparative genetic characterization of Enteroaggregative *Escherichia coli* strains recovered from clinical and non-clinical settings. Sci Rep. 2016;6:24321.doi:10.1038/srep24321

74. Foster-Nyarko E, Alikhan NF, Ikumapayi UN, Sarwar G, Okoi C, Tientcheu PM, et al. Genomic diversity of *Escherichia coli* from healthy children in rural Gambia. PeerJ. 2021;9, e10572. 10.7717/peerj.10572

75. Zueter AM, Mharib T, Shqair D, Al-Tamimi M, Sawan HM, Zaiter A, et al. Multi-locus sequence typing of Escherichia coli isolated from clinical samples in Jordan. J Infect Dev Ctries. 2024;18(4):571–578. doi: 10.3855/jidc.18810.

76. Hoshiko Y, Chowdhury G, Kitahara K, Ghosh D, Nagano DS, Ohno A, et al. Genomic features of three major diarrhoeagenic *Escherichia coli* pathotypes in India. Microb Genom. 2025;11(7):001430. doi: 10.1099/mgen.0.001430.

77. Aworh MK, Kwaga JKP, Hendriksen RS, Okolocha EC, Thakur S. Genetic relatedness of multidrug resistant Escherichia coli isolated from humans, chickens and poultry environments. Antimicrob Resist Infect Control. 2021;10(1):58. doi: 10.1186/s13756-021-00930-x.

78. Dias RCB, Tanabe RHS, Vieira MA, Cergole-Novella MC, Dos Santos LF, Gomes TAT, et al. Analysis of the Virulence Profile and Phenotypic Features of Typical and Atypical Enteroaggregative *Escherichia coli* (EAEC) Isolated From Diarrheal Patients in Brazil. Front Cell Infect Microbiol. 2020;10:144. doi: 10.3389/fcimb.2020.00144.

79. Bhargava S, Johnson BB, Hwang J, Harris TA, George AS, Muir A, et al. Heat-resistant agglutinin 1 is an accessory enteroaggregative Escherichia coli colonization factor. J Bacteriol. 2009;191(15):4934–42. doi: 10.1128/JB.01831-08.

80. Tiwari SK, van der Putten BCL, Fuchs TM, Vinh TN, Bootsma M, Oldenkamp R, et al. Genome-wide association reveals host-specific genomic traits in Escherichia coli. BMC Biol. 2023;21(1):76. doi: 10.1186/s12915-023-01562-w.

81. Wang J, Lu Q, Yao L, Zhang W, Hu Q, Guo Y, et al. Research note: Prevalence and genetic characteristics of pathogenic E. coli isolates from domestic pigeons in central China. Poult Sci. 2025;104(2):104705. doi: 10.1016/j.psj.2024.104705.

82. Yang GY, Guo L, Su JH, Zhu YH, Jiao LG, Wang JF. Frequency of Diarrheagenic Virulence Genes and Characteristics in *Escherichia coli* Isolates from Pigs with Diarrhea in China. Microorganisms. 2019;7(9):308. doi: 10.3390/microorganisms7090308.

83. Kaper JB, Nataro JP, Mobley HL. Pathogenic *Escherichia coli*. Nat Rev Microbiol. 2004;2(2):123–40. doi: 10.1038/nrmicro818.

