## Supplementary material for "Enteroaggregative *Escherichia* clade I from Nigeria": Suppl Tables 1 to 4

**ENTEROAGGREGATIVE NON-COLI SUPPLEMENTARY MATERIALS**

**Supplementary Table 1:** Phylogenetic group and Microbact ID of *coli* and non-*coli Escherichia*

| **Name** | **Genome type** | **PhyloGrp** | **MicrobaOct** | **MicrobaID** | **Phylogeny** |
| --- | --- | --- | --- | --- | --- |
| A98YA | case | F |  |  | *E.* Clade I |
| LLH131E | control | F | 27600461 | *E. coli* | *E.* Clade I |
| LWD014D | case | F | 47600660 | *E. coli* | *E.* Clade I |
| LWD014E | case | F | 47600660 | *E. coli* | *E.* Clade I |
| LWD014G | case | F | 67600360 | *E. hermannii* | *E.* Clade I |
| LWD014H | case | F | 66600460 | *E. coli* | *E.* Clade I |
| MND079F | case | ~~F~~ |  |  | *E.* Clade I |
| MND79B | case | F |  |  | *E.* Clade I |
| LLH131E2 | control | F | 27600461 | *E. coli* | *E.* Clade I |
| E16 | case | F |  |  | *E.* Clade I |
| E07 | case | G |  |  | *E.* Clade I |
| LLH235F | control | U/C |  |  | *E. fergusonii* |
| LLH071B | control | U/C | 67600061 | *E. coli* | *E. ruysiae* |
| LLH071E | control | U/C | 66600040 | *E. coli* | *E. ruysiae* |
| CHD059C | case | A | 67600764 | *E. coli* | *E. coli* |
| CHD016E | case | A | 66620620 | *E. coli* Inactive | *E. coli* |
| LKH141A | control | A | 07600461 | *E. coli* Inactive | *E. coli* |
| LKH209F | control | A | 47600621 | *E. coli* Inactive | *E. coli* |
| LLD021C | case | A | 46600660 | *E. coli* | *E. coli* |
| CHD041E | case | B1 | 67600760 | *E. coli* | *E. coli* |
| CHD086D | case | B1 | 67600660 | *E. coli* | *E. coli* |
| LLH005D | control | B1 | 6770006 | *E. coli* | *E. coli* |
| LLH070E | control | B1 | 67600266 | *E. hermannii* | *E. coli* |
| LLD019A | case | B1 | 67600664 | *E. coli* | *E. coli* |
| LKD69I | case | B2 | 67600720 | *E. coli* | *E. coli* |
| LLH021A | control | B2 | 66700721 | *E. coli* | *E. coli* |
| CHD069C | case | B2 | 67600764 | *E. coli* | *E. coli* |
| LWD023E | case | B2 | 67600660 | *E. coli* | *E. coli* |
| MNH136H | control | B2 | 67200621 | *E. coli* Inactive | *E. coli* |
| CHD076F | case | C | 66600660 | *E. coli* | *E. coli* |
| CHD076I | case | C | 47600060 | *E. coli* Inactive | *E. coli* |
| CHD076J | case | C | 67600260 | *E. hermannii* | *E. coli* |
| LKH215F | control | C | 47600222 | *E. coli* Inactive | *E. coli* |
| LKH215H | control | C | 47600700 | *E. coli* Inactive | *E. coli* |
| CHD020E | case | D | 47600620 | *E. coli* | *E. coli* |
| CHD063A | case | D | 47600464 | *E*. *coli* | *E. coli* |
| LLH201B | control | D | 67600461 | *E. coli* | *E. coli* |
| LLD53H | case | D | 47600261 | *E. coli* | *E. coli* |
| LLH191C | control | D | 67763777 | *K. ornithinolytica* | *E. coli* |
| LLH150B | control | E | 47600661 | *E. coli* | *E. coli* |
| LLH198E | control | E | 00540000 | *A. haemolyticus* | *E. coli* |
| CHD086A | case | E | 67600740 | *E. coli* | *E. coli* |
| LKH209I | control | F | 47600420 | *E. coli* Inactive | *E. coli* |
| LKH209J | control | F | 46600420 | *E. coli* Inactive | *E. coli* |
| LLH054C | control | F | 66600420 | *E. coli* Inactive | *E. coli* |
| CHD071A | case | F | 67600560 | *E. coli* | *E. coli* |
| CHD033C | case | G | 67600660 | *E. coli* | *E. coli* |
| JKH044E | control | G | 67600760 | *E. coli* | *E. coli* |
| LLH051A | control | G | 47600660 | *E. coli* | *E. coli* |
| LLH060C | control | G | 67600660 | *E. coli* | *E. coli* |
| LLH099D | control | G | 47700060 | *E. coli* | *E. coli* |
| EAEC 042 | case | D |  | *E. coli* | *E. coli* |

U/C – undefined/cryptic phylogenetic group

**Supplementary Table 2:** Core SNP differences between putative non-*coli Escherichia* and reference genomes

| **Strain** | **Nearest reference genome** | **SNP differences from nearest reference** | **SNP differences from *E*. *coli* reference (EAEC 042)** | **ID (Phylogeny)** |
| --- | --- | --- | --- | --- |
| A98YA | ERR1790650 | 415 | 129671 | *E.* Clade I |
| LLH131E | SRR35332963 | 180 | 57121 | *E.* Clade I |
| LWD014D | SRR11848721 | 659 | 128068 | *E.* Clade I |
| LWD014E | SRR12495206 | 379 | 130337 | *E.* Clade I |
| LWD014G | SRR12495206 | 323 | 129786 | *E.* Clade I |
| LWD014H | SRR12495206 | 306 | 132638 | *E.* Clade I |
| MND079F | SRR11849145 | 607 | 130163 | *E.* Clade I |
| MND79B | SRR11848721 | 662 | 130484 | *E.* Clade I |
| LLH131E2 | SRR11848858 | 339 | 130104 | *E.* Clade I |
| E16 | ASM3020798v1 | 636 | 52067 | *E.* Clade I |
| E07 | ERR1790650 | 339 | 118646 | *E.* Clade I |
| LLH235F | SRR17304136 | 29060 | 140834 | *E. fergusonii* |
| LLH071B | SRR17304136 | 29455 | 27699 | *E. ruysiae* |
| LLH071E | SRR17304136 | 29454 | 221517 | *E. ruysiae* |
| CHD274C | SRR17304136 | *28437* | 103070 | *E. coli* |
| CHD277D | SRR17304136 | *28115* | 88254 | *E. coli* |
| CHD287B | SRR17304136 | *28084* | 84670 | *E. coli* |
| CHD313B | SRR17304136 | 27887 | 71343 | *E. coli* |
| CHD338E | SRR17786805 | *26955* | 87015 | *E. coli* |
| CHH130E | SRR17786805 | *11745* | 67536 | *E. coli* |
| CHH211D | SRR17786805 | *25351* | 91797 | *E. coli* |
| JKH044G | SRR17304136 | *28060* | 87892 | *E. coli* |
| LKH209J | SRR17304136 | *28087* | 86668 | *E. coli* |
| LLD091B | SRR17304136 | *28047* | 71603 | *E. coli* |
| LLD09B | SRR17304136 | 28383 | 103826 | *E. coli* |
| LLD108A | SRR17786805 | 25764 | 93867 | *E. coli* |
| LLD52D1 | SRR17304136 | 28267 | 98248 | *E. coli* |
| LLH060C | SRR17304136 | 28112 | 83124 | *E. coli* |
| LLH070E | SRR17786805 | 23490 | 90783 | *E. coli* |
| LLH10E | SRR17304136 | 28127 | 85114 | *E. coli* |
| LLH120G | SRR17304136 | *28159* | 90487 | *E. coli* |
| LLH19A | SRR17304136 | *28292* | 101253 | *E. coli* |
| LLH238A | SRR17304136 | 28073 | 83528 | *E. coli* |
| LLH249H | SRR17304136 | 28248 | 96947 | *E. coli* |
| LLH45E | SRR17304136 | 28024 | 61109 | *E. coli* |
| LWD45B | SRR17786805 | 24823 | 92494 | *E. coli* |
| MND095D | SRR17304136 | 28248 | 97679 | *E. coli* |
| MNH314E | SRR17786805 | 25138 | 92842 | *E. coli* |
| EAEC 042 | SRR17304136 | 27699 | 0 | *E. coli* |

**Supplementary Table 3**: Summary of ST5792 Genomes found on Enterobase

| **S/No** | **Accession Number** | **Source of Isolation** | **Continent** | **Country** | **Year of isolation** | **Species (Enterobase)** | **ST** |
| --- | --- | --- | --- | --- | --- | --- | --- |
| **1.** | SRR1186783 | Human | Africa | Tanzania | 2009 | Clade IV | 5792 |
| **2.** | ERR1640608 | NA | NA | NA | NA | Clade IV | 5792 |
| **3.** | SRR7693687 | Human | NA | NA | 2014 | Clade IV | 5792 |
| **4.** | SRR9887404 | Environment | Europe | Sweden | 2013 | Clade IV | 5792 |
| **5.** | NA | NA | NA | NA | NA | NA | 5792 |
| **6.** | NA | NA | NA | NA | NA | NA | 5792 |
| **7.** | NA | NA | NA | NA | NA | NA | 5792 |
| **8.** | SRR29019528 | Human | Asia | Bangladesh | 2019 | *E. coli* | 5792 |
| **9.** | SRR29019529 | Human | Asia | Bangladesh | 2019 | *E. coli* | 5792 |
| **10.** | NA | NA | NA | NA | NA | NA | 5792 |
| **11.** | NA | NA | NA | NA | NA | NA | 5792 |

**Supplementary Table 4:** Summary of ST5636 Genomes found on Enterobase

| **S/No** | **Accession Number** | **Source of Isolation** | **Continent** | **Country** | **Year of isolation** | **Species (Enterobase)** | **ST** |
| --- | --- | --- | --- | --- | --- | --- | --- |
| **1.** | SRR1916750 | Food | North America | Canada | 2014 | *E. fergusonii* | ST5636 |
| **2.** | NA | NA | Asia | China | 2020 | NA | ST5636 |
| **3.** | NA | NA | Asia | China | 2020 | NA | ST5636 |
| **4.** | NA | NA | Asia | China | 2020 | NA | ST5636 |
| **5.** | NA | NA | Asia | China | 2021 | NA | ST5636 |
| **6.** | NA | NA | Asia | China | 2021 | NA | ST5636 |
| **7.** | NA | NA | Asia | China | 2021 | NA | ST5636 |
| **8.** | NA | NA | Asia | China | 2022 | NA | ST5636 |
| **9.** | NA | NA | Asia | China | 2022 | NA | ST5636 |
| **10.** | NA | NA | Asia | China | 2022 | NA | ST5636 |
| **11.** | NA | NA | Asia | China | 2022 | NA | ST5636 |
| **12.** | ASM1382210v1 | NA | Europe | UK | NA | NA | ST5636 |
| **13.** | ASM2601229v1 | NA | Asia | Japan | NA | NA | ST5636 |
| **14.** | ASM2601231v1 | NA | Asia | Japan | NA | NA | ST5636 |
| **15.** | NA | NA | Asia | China | 2019 | NA | ST5636 |
| **16.** | ASM1389243v1 | NA | Europe | UK | NA | NA | ST5636 |
| **17.** | NA | NA | Asia | China | 2014 | NA | ST5636 |
| **18.** | NA | NA | Asia | China | 2017 | NA | ST5636 |
| **19.** | NA | NA | Asia | China | 2017 | NA | ST5636 |
| **20.** | NA | NA | Asia | China | 2020 | NA | ST5636 |
| **21.** | NA | NA | Asia | China | 2020 | NA | ST5636 |
